# Computing Stackelberg Equilibrium for Cancer Treatment

**DOI:** 10.1101/2024.09.09.612059

**Authors:** Sam Ganzfried

## Abstract

Recent work by Kleshnina et al. has presented a Stackelberg evolutionary game model in which the Stackelberg equilibrium strategy for the leading player corresponds to the optimal cancer treatment [4]. We present an approach that is able to quickly and accurately solve the model presented in that work.

## 1 Introduction and problem formulation

In this section we review the Stackelberg evolutionary game dynamic model of cancer evolution previously studied [4]. There are two players: a follower and a leader. The leader is a physician who selects amounts of two different drugs to use for therapy, *m*_1_ and *m*_2_. The follower is a cancer population consisting of three cell types: 0-type denotes a cell that does not develop resistance to either drug, 1-type cells are resistant to just drug 1, and 2-type cells are resistant to just drug 2. The follower selects a population size for each type, denoted *x*_0_, *x*_1_, *x*_2_, as well as a trait for each type, denoted *u*_0_, *u*_1_, *u*_2_. It is assumed that each of these variables are nonnegative, with the *u*_*i*_ ∈ [0, 1]. It is also assumed that they are all implicitly functions of time *t*. Note that *u*_0_ does not appear anywhere in the analysis so can be ignored.

For each cell type *i* there is a fitness function *G*_*i*_(*u*_*i*_, **m, x**) that the follower is trying to maximize. We assume that the dynamics of the population *x* are governed by

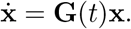

In order to ensure that we are in equilibrium of the ecological dynamics we must have that 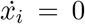 for *i* = 0, 1, 2. Thus, the follower is selecting *u*_1_ that maximizes *G*_1_, *u*_2_ that maximizes *G*_2_, and *x*_0_, *x*_1_, *x*_2_ that ensure equilibrium of the ecological dynamics. The leader, knowing that the follower will subsequently select their actions in this way, selects *m*_1_, *m*_2_ to maximize a quality of life function *Q*(**m, u, x**).

Thus, we can formulate the problem of determining the optimal strategies for both players as follows:

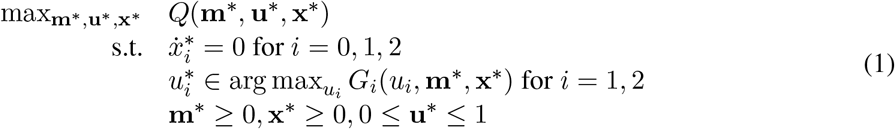

This general model is instantiated by the following functional forms for the fitness functions *G*_0_, *G*_1_, *G*_2_ and quality of life function *Q*. Note that the model presentation is slightly different between the paper [4] and the implementation in the code repository [3]. We will be using the model presented in the code.

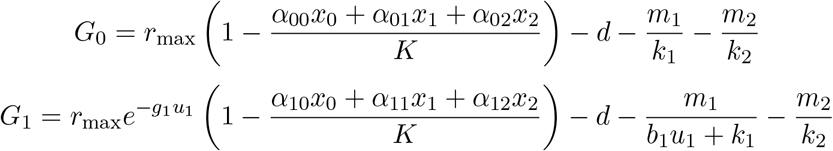

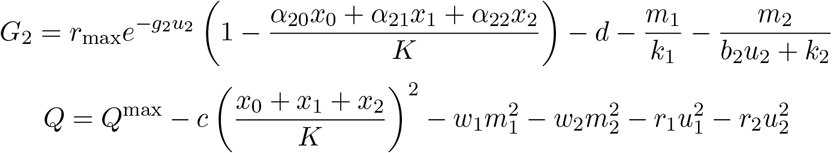

The model has several parameters, whose interpretations are summarized in Table 1. Note that in the code additional parameters *a*_0_, *a*_1_, *a*_2_, *a*_3_ are defined, with

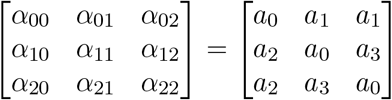

**Table 1:** Interpretations of model parameters.

| Parameter | Interpretation |
| --- | --- |
| $r_{\max}$ | Max cell growth rate |
| $g_i$ | Cost of resistance strategy (cell type) $i$ |
| $\alpha_{ij}$ | Interaction coefficient between cell types $i$ and $j$ |
| $K$ | Carrying capacity |
| $d$ | Natural death rate |
| $k_i$ | Innate resistance that may be present before drug exposure |
| $b_i$ | Benefit of the evolved resistance trait in reducing therapy efficacy |
| $Q^{\max}$ | Quality of life of a healthy patient |
| $w_i$ | Toxicity of drug $i$ |
| $r_i$ | Effect of resistance rate of cell type $i$ |
| $c$ | Weight for impact of tumor burden vs. drug toxicity/treatment-induced resistance rate |

## 2 Prior approach

In this section we present the approach described in Github repository created by the authors [3]. They first solve for expressions for *x*_*i*_ in terms of *u*_*i*_, *m*_*i*_ such that the condition for equilibrium of the ecological dynamics is satisfied. This involves solving a system of three equations *G*_*i*_*x*_*i*_ = 0 with unknowns *x*_1_, *x*_2_, *x*_3_. They calculate the following analytical solution:

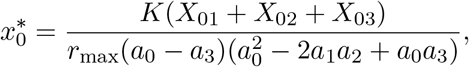

where *X*_01_, *X*_02_, *X*_03_ denote the following quantities:

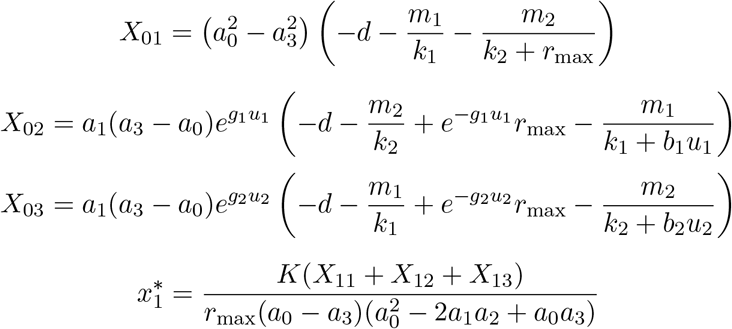

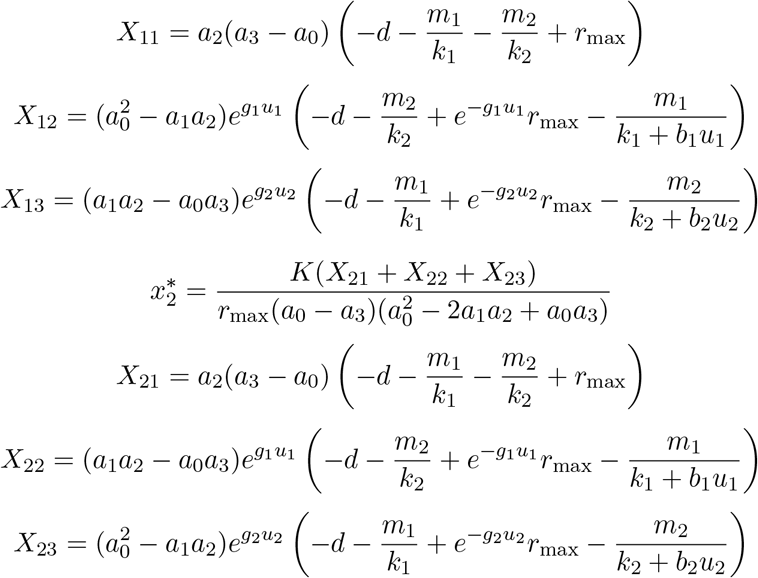

While it is not given in the model formulation, the code assumes that *m*_1_ and *m*_2_ fall in the interval 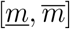, with 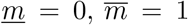. They discretize the space so that *m*_1_ and *m*_2_ are multiples of *h* = 0.1, and iterate over all combinations, of which there are 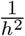. For each combination of values they calculate the optimal values of *u*_1_, *u*_2_ as follows. They assume that the optimal value of *u*_1_ is where 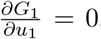, and the optimal value of *u*_2_ is where 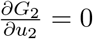. They calculate the following expressions for the partial derivatives:

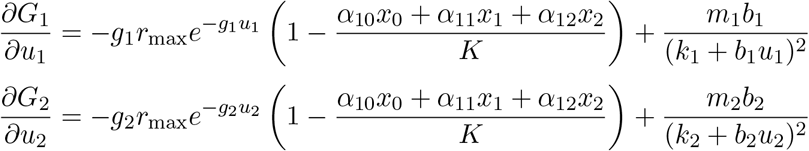

To find values of *u_1_, u*_2_ for which these derivatives equal zero, they perform an optimization to minimize the quantity 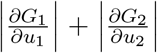. They perform this optimization in Matlab using the fminsearchbnd function, using bounds 0 ≤*u*_1_ ≤1, 0 ≤*u*_2_ ≤1. Note that the above expressions for 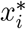 which are in terms of *m*_*i*_ and *u*_*i*_ are substituted into the expressions for 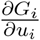, so that *u*_*i*_ is the only variable in the objective. This procedure results in optimal values 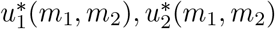 for each combination of values for (*m*_1_, *m*_2_). The values for 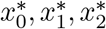 are determined by these values.

Finally, they iterate over all 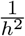 of these combinations of values to determine which one maximizes the value of *Q*. This determines the optimal values of 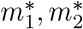 for the leader, which in turn determine the optimal strategies for the follower. The overall procedure is summarized in Algorithm 1.

### Algorithm 1

Prior approach

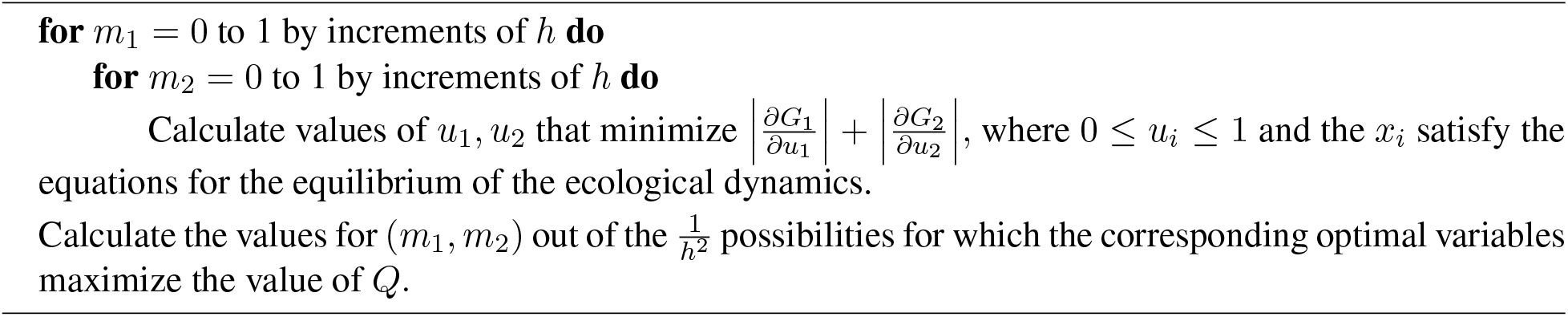

## 3 Limitations of prior approach

The prior approach has several significant limitations. The first is that it does not check that the calculated values for 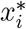 for given values of *u*_*i*_ and *m*_*i*_ are biologically sensible. Since the *x*_*i*_ correspond to populations, they must be nonnegative.

Another limitation is that the coarse discretization of values for *m*_*i*_ means that only a small number of possibilities are considered. This also means that the running time will potentially be large, since we must perform 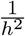 separate optimizations.

Another significant limitation is that it is assumed that *G*_*i*_ is maximized when 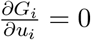, and the boundary cases when it is maximized at *u*_*i*_ = 0 or 1 are ignored.

A final limitation is that the procedure invokes the fminsearch algorithm in Matlab, which is not even guaranteed to find a local minimum, let alone a global minimum.

## 4 New approach

We now describe our new approach that addresses the limitations of the prior approach. We will formulate a single quadratic program that corresponds to the full optimization problem and solve it using Gurobi’s nonconvex MIQCP solver which has a guarantee of global optimality (subject to numerical precision).

First we have the main decision variables *x*_*i*_, *u*_*i*_, *m*_*i*_ with *x*_*i*_ ≥0, *m*_*i*_ ≥0, 0 ≤*u*_*i*_ ≤1. The objective function *Q* is a quadratic function of these variables. Next we encode the conditions for equilibrium of the ecological dynamics. We must define several auxiliary variables to do this.

First define *η*_*i*_ = *g*_*i*_*u*_*i*_, and 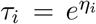 for *i* = 1, 2. For the latter, we use Gurobi’s addGenConstrExp function that uses a piecewise linear approximation for the exponential function. We set these variables to be nonnegative. To provide tighter upper bounds we can set *η*_*i*_ ≤*g*_*i*_, 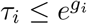, since *u*_*i*_ 1. We next define the auxiliary variable 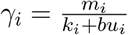. We can do this by including the quadratic constraint *k*_*i*_*γ*_*i*_+*bu*_*i*_*γ*_*i*_ −*m*_*i*_ = 0 for *i* = 1, 2. Using these variables we can now encode the conditions for equilibrium of ecological dynamics using constraints that are quadratic in the variables.

Next we must encode the conditions that *u*_*i*_ is a maximizer of *G*_*i*_. To do this we define several additional auxiliary variables. We define 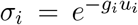 by adding in the constraint *σ*_*i*_*τ*_*i*_ = 1. Next we define 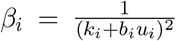. We can do this by including the quadratic constraint 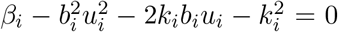. Finally we define 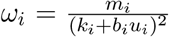 by including the quadratic constraint *ω*_*i*_*β*_*i*_ − *m*_*i*_ = 0. Using these variables, we can now encode the expressions for 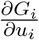 that are quadratic in the variables.

Recall that we are trying to select *u*_*i*_ ∈ [0, 1] to maximize *G*_*i*_, for *i* = 1, 2. We can do this by introducing two Lagrange multipliers *λ*_*i*1_ ≥0, *λ*_*i*2_ ≥0. Then the KKT optimality condition is equivalent to the following three constraints:

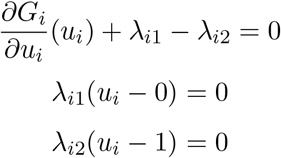

These constraints are all quadratic in the variables and ensure that we find *u*_*i*_ ∈ [0, 1] that maximizes *G*_*i*_ regardless of whether it is at the boundary or at an interior solution with the derivative equal to zero.

Our full formulation is given below. Here the *X*_*ij*_ correspond to the same quantities as before and are just defined to simplify presentation, not as new variables.

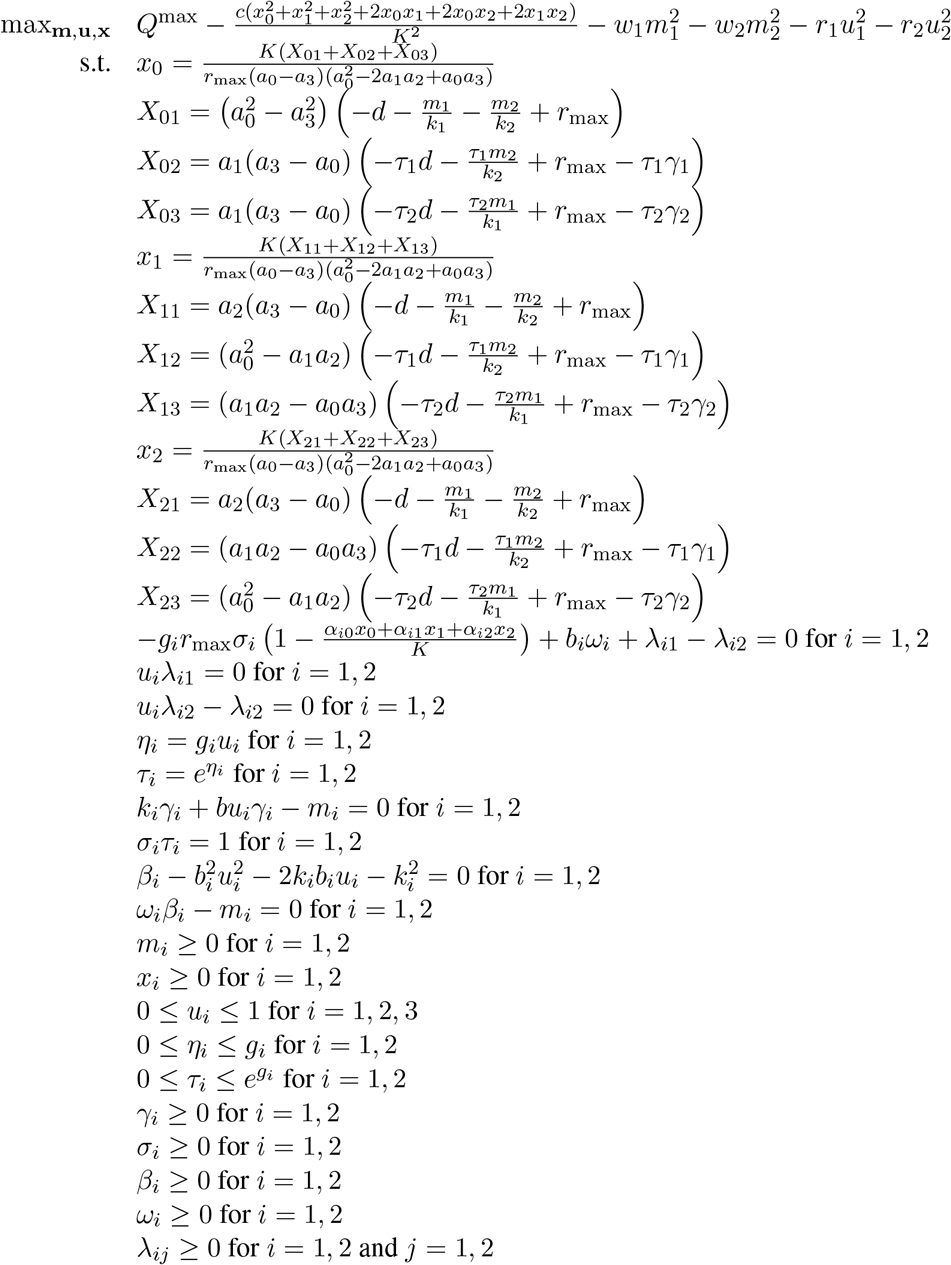

This formulation addresses the limitations of the prior approach. It ensures that all quantities are biologically relevant by imposing nonnegativity constraints on corresponding variables. It allows *m*_*i*_ to take on arbitrary nonnegative values, not a small set of discretized values. It involves solving a single optimization problem instead of 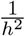 separate optimization problems. It uses KKT conditions to ensure that the values of *u*_*i*_ that maximize *G*_*i*_ are found regardless of whether they are interior or boundary solutions. And the approach guarantees finding a global optimum since that is guaranteed by Gurobi’s nonconvex MIQCP solver.

## 5 Experiments

We ran experiments with both approaches on a problem instance using the same parameter values as the prior approach [3], which are provided in Table 2. All experiments were done on a single core of a laptop using Windows 11. Experiments with the prior approach used Matlab version 24.1.0.2689473 (R2024a) Update 6 [2], and experiments with the new approach were done using Gurobi version 11.03 [1] with Java version 14.0.2. For the optimizations in the prior approach, Matlab’s function fminsearchbnd was called using parameters TolX = 1 *×* 10^−12^, MaxFunEvals = 1000. The results are shown in Table 3. We can see that the prior approach found a solution with a negative value for 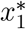, which is not biologically sensible. The prior approach took nearly five minutes while our new approach took less than two seconds.

**Table 2:** Parameter values used in experiments.

| Parameter | Value |
| --- | --- |
| $r_{\max}$ | 0.45 |
| $g_1$ | 0.5 |
| $g_2$ | 0.5 |
| $a_0$ | 1 |
| $a_1$ | 0.15 |
| $a_2$ | 0.9 |
| $a_3$ | 0.9 |
| $K$ | 10,000 |
| $d$ | 0.01 |
| $k_1$ | 5 |
| $k_2$ | 5 |
| $b_1$ | 10 |
| $b_2$ | 10 |
| $Q^{\max}$ | 1 |
| $w_1$ | 0.5 |
| $w_2$ | 0.2 |
| $r_1$ | 0.4 |
| $r_2$ | 0.4 |
| $c$ | 0.5 |

**Table 3:** Experimental results for both approaches.

|  | Prior approach | New approach |
| --- | --- | --- |
| $m_1^*$ | 0.4 | 0.40837 |
| $m_2^*$ | 0.5 | 0.46579 |
| $u_1^*$ | 0.19015 | 0.21361 |
| $u_2^*$ | 0.3123 | 0.28554 |
| $x_0^*$ | 5634.3774 | 5749.8474 |
| $x_1^*$ | -360.2658 | 1.3366 |
| $x_2^*$ | 1316.2683 | 950.5000 |
| $Q^*$ | 0.59936 | 0.59780 |
| Running time (seconds) | 282.69 | 1.65 |

## 6 Conclusion

We presented a new approach for computing Stackelberg equilibrium strategies in a Stackelberg evolutionary game dynamic model of cancer evolution previously studied. Our approach is based on solving a new quadratic program formulation. We noted several limitations of the approach used by prior work which are addressed by our approach. When we compared the approaches on a sample instance our approach ran significantly faster and the prior approach output a solution that is not biologically relevant. As more complex game-theoretic and optimization models are being formulated for problems in biology and cancer treatment in particular, it is important to develop efficient algorithms for accurately solving them. While we focused on one instantiation presented in prior work, our approach is applicable more generally to computing optimal strategies in Stackelberg evolutionary games.

